# Semantic distance differently modulates FPVS-EEG responses to words and pictures

**DOI:** 10.64898/2026.02.26.706120

**Authors:** Angelique Volfart, Aliette Lochy, Bruno Rossion, Matthew A. Lambon Ralph

**Author notes:** **Corresponding author:** Dr. Angelique Volfart, Université du Luxembourg, Maison des Sciences Humaines 11 Porte des Sciences, 4366 Belval Esch-sur-Alzette, Luxembourg. shared last authorship (Angelique Volfart, Bruno Rossion, Matthew A. Lambon Ralph).

## Abstract

The organization of semantic knowledge within the brain has long been studied through various theoretical frameworks and is still a matter of debate. Many studies that have helped advancing our knowledge about this topic have approached it in the context of its deterioration, notably in patients with semantic dementia. In healthy subjects, the question whether and how semantic distance between concepts represented by picture or word stimuli influences brain responses remains understudied. In this study with 24 healthy subjects, we used electroencephalography (EEG) recordings coupled with a fast periodic visual stimulation (FPVS) approach, for the first time, to assess the semantic distance effect with different stimulus modalities. Picture or word stimuli from a reference category (birds) appeared every fourth item among a 4-Hz stream of either man-made objects (forming the high distance [HD] condition) or other animals (low distance [LD] condition). Within a few minutes of recording, EEG responses were observed at the pre-determined frequency of the reference category presentation (at 1 Hz), suggesting its exemplars were activated at least sufficiently to be automatically discriminated from different superordinate (HD) and basic (LD) categories, for both pictures and words. Pictures and words elicited a different pattern of response to semantic distance, with larger amplitudes for HD than LD conditions for pictures but a reversed pattern of somewhat greater amplitudes for LD than HD conditions for words. Our findings support a similarity-based structure of semantic representations and provide evidence for a differential mapping between semantic and picture/word surface representations.

## 1 Introduction

Semantic memory is a core aspect of human cognition, subtending our general knowledge of words, objects, people, etc. The cognitive and anatomical organization of semantic knowledge has long been studied through various theoretical frameworks and is still a matter of debate (see Frisby et al., 2023; Lambon Ralph et al., 2017 for reviews).

The hub-and-spoke model provides a powerful computational framework for understanding how concepts are encoded and differentiated in the brain. This model, supported by neuropsychological and computational studies (e.g., Coccia et al., 2004; Lambon Ralph & Howard, 2000; Patterson et al., 2007; T. T. Rogers et al., 2004, 2015; T. T. Rogers & Patterson, 2007; see also Lambon Ralph et al., 2017), suggests that semantic representations are distilled from our multimodal experiences, with the resultant representational geometry reflecting the graded similarity structure between concepts (T. T. Rogers et al., 2004; T. T. Rogers & McClelland, 2004). Its neural basis has been hypothesized to be centred in the anterior part of the temporal lobes, a region densely interconnected with sensory and association cortices and thus ideally located for fostering communication with multiple modality-specific regions distributed across the brain (Bajada et al., 2019; e.g., Binney et al., 2012). The role of the anterior temporal lobes in representing semantic knowledge has been supported by many studies with diverse methodological approaches (e.g., lesion studies: Binney et al., 2010; Hoffman et al., 2014; Rice et al., 2018; neuroimaging: Hoffman & Lambon Ralph, 2018; Rice et al., 2015; Visser et al., 2010; brain stimulation: Lambon Ralph et al., 2009; Pobric et al., 2010; Shimotake et al., 2015; electrophysiology: Chen et al., 2016; Jackson et al., 2015; T. T. Rogers et al., 2021).

In the computational implementation of this model, semantic representations are conceived as activation patterns across an ensemble of points in a multidimensional vector space, where conceptual similarity/dissimilarity depends upon the distance between different points within this space (Cox et al., 2024; Hoffman et al., 2018; Jackson et al., 2021). These semantic representations are “transmodal”, i.e., not tied to any individual perceptual modality, which means they cannot be inferred from activation patterns elicited by one or another modality-specific surface representation alone (T. T. Rogers et al., 2004). This is particularly important because each modality structures concepts in distinct ways. For example, based on visual information, concepts will be structured according to their similarity in shape, colour or size, while, based on functional information, they will be organized according to one’s experience of interaction and use of them (Dilkina & Lambon Ralph, 2013; Fernandino et al., 2022), requiring the semantic system to abstract from these and built its own, transmodal similarity structure. These semantic representations provide the basis for generalization and inference, i.e., the ability to generalize across concepts that are semantically similar despite being perceptually different, and to infer attributes of newly encountered objects based on their conceptual similarity with previously stored knowledge (T. T. Rogers et al., 2004; see also Lambon Ralph et al., 2010; Lambon Ralph & Patterson, 2008).

While many studies have tried to understand the organization of conceptual categories based on pictures (e.g., Connolly et al., 2012; Devereux et al., 2018), assessing the hub-and-spoke’s proposal of similarity-based structure requires probing the system with different stimulus modalities (e.g., words). An important point of this framework notably relates to the differential mapping assumed between conceptual and different modality-specific surface representations (e.g., pictures and words). Indeed, several neuropsychological studies in patients with semantic dementia (SD), a disorder specifically affecting semantic memory (at least in the early stages of the disease), have reported a differential pattern of performance when accessing semantic representations from pictures or words, with usually worse performance with words than pictures (e.g., Lambon Ralph & Howard, 2000; T. T. Rogers et al., 2004; see also Coccia et al., 2004). To account for these behavioural differences, the model implements a differential mapping between conceptual knowledge and surface representations of words vs. pictures: while pictures have some systematic relationships with conceptual representations, words adopt an arbitrary mapping with the latter (T. T. Rogers et al., 2004; see also Caramazza et al., 1990). Computational simulations of such model (Lambon Ralph & Howard, 2000; T. T. Rogers et al., 2004) show that it can indeed lead to worse performance for words than pictures when the semantic representations are damaged, as well as faster degradation of performance for words with increasing damage.

Overall, the similarity-based semantic structure proposed by the hub-and-spoke framework has been demonstrated to explain many behavioural patterns observed under both undamaged and damaged conditions, with most supporting evidence coming from SD patients (e.g., T. T. Rogers et al., 2004; T. T. Rogers & Patterson, 2007). However, less has been done to confirm this structure in the healthy brain.

The fast periodic visual stimulation (FPVS) approach coupled with electroencephalography (EEG) is a powerful tool to offer a window into the healthy brain’s semantic organization. With this approach, objects belonging to a given semantic category can be visually presented at a fast frequency rate (i.e., base stimuli at 4 Hz) while objects belonging to another category appear every 4 items (i.e., alternate stimuli at 1 Hz), and participant’s electrophysiological responses at these pre-determined frequencies are analysed (e.g., Volfart et al., 2021). In order to generate an “odd-one-out” FPVS response at 1 Hz, the underlying neurocognitive process must have differentiated between the two categories being presented. In addition to being a direct measure of this discrimination process, the FPVS approach has multiple benefits, allowing the recording of objective, high-SNR responses at the frequency bin of interest, in a short amount of time and without the need for an explicit semantic task.

In the present study, we investigated the similarity structure of the semantic space with pictures and written words with a paradigm in which object concepts belonging to a reference category (birds) were presented every fourth item in a 4-Hz stream of items from various semantic categories (instead of only one category, as previously done in David et al., 2025; Volfart et al., 2021). Reference stimuli presented at 1 Hz were always exemplars of the bird category (e.g., swan, owl, etc.) and two conditions were formed based on the semantic distance between this reference category and the base stimuli. In the high semantic distance (HD) condition, base stimuli all belonged to a different superordinate category (i.e., man-made) but could be drawn from various basic categories (e.g., musical instruments, vehicles, etc.), while in the low semantic distance (LD) condition, base stimuli belonged to the same superordinate category as the reference stimuli (i.e., animals) but were drawn from distinct basic categories (e.g., rodents, felines, etc.).

Using this FPVS-EEG approach, we addressed two questions. First, are word and picture stimuli displaying exemplars of the reference category automatically and implicitly discriminated from exemplars of other categories? Based on previous FPVS studies (e.g., Stothart et al., 2017; Volfart et al., 2021), we expected to observe EEG responses at 1 Hz reflecting the discrimination of base and reference stimuli in both the HD and LD conditions, because reference stimuli all belong to the same basic category of birds while base stimuli all belong either to a different superordinate category (man-made) or to other basic categories of animals (e.g., rodents, dogs). If EEG discrimination responses are observed, are these different for pictures and words? We hypothesized that EEG discriminative responses to picture conditions would have larger amplitudes than responses to word conditions, regardless of the semantic distance condition, due to the more systematic mapping between conceptual knowledge and surface form for pictures than words.

Second, could we observe a semantic distance effect in EEG discrimination responses, i.e., differential responses whether items in the base category are semantically closer or further away from items in the reference category? If there is a semantic distance effect in EEG responses, does it differ for pictures and words? We hypothesized that: a) there will be a strong semantic distance effect for pictures (as manifested by differential amplitudes in HD vs LD conditions), in the direction of larger amplitudes in the HD condition considering the likely greater contribution of visual features for distinguishing stimuli in the HD vs. LD contrasts (i.e., discriminating objects across superordinate or within basic categories; Ashtiani et al., 2017; Collin & Mcmullen, 2005; Praß et al., 2013) and previous findings of better performance with superordinate categories at rapid stimulus presentation rates (e.g., Fabre-Thorpe, 2011; Macé et al., 2009; Praß et al., 2013; T. T. Rogers & Patterson, 2007); b) the semantic distance effect will be weaker for words than pictures, based on SD patient studies and computational explorations showing that performance in semantic tasks drops off more quickly for words than pictures (e.g., Lambon Ralph & Howard, 2000; T. T. Rogers et al., 2004).

## 2 Materials and Methods

### 2.1 Participants

Twenty-five participants were recruited among students at the Queensland University of Technology (Brisbane, Australia). One participant was later excluded due to technical issues during EEG recording, resulting in a final sample of 24 participants (mean age = 21.6 ± 3.8; 15 females). This sample size was determined *a priori* based on previous FPVS-EEG studies using the same approach with samples of 10-20 participants (n = 22: Lochy et al., 2024; e.g., n = 20: Stothart et al., 2017; n = 14: Volfart et al., 2021). In addition, we conducted an a priori power analysis in GPower (version 3.1.9.7) for our main repeated measures ANOVA, which tested the interaction between Item Type (Pictures vs. Words), Condition Type (Control vs. Experimental), and Distance (HD vs. LD). Using Cohen’s f = 0.51 as a proxy from a similar study by Volfart et al. (2021), correlation among repeated measures r = 0.46 (estimated from EEG amplitude data), and non-sphericity correction ε = 0.75, GPower indicated that only n ≈ 6 participants would be sufficient to achieve 80% power at α = .05. Our final sample of 24 participants therefore substantially exceeds this requirement, ensuring robust power even under more conservative assumptions.

All participants were right-handed, native English speakers with normal or corrected-to-normal vision and normal hearing. Participants reported no history of neurological disorders, psychiatric conditions, learning disabilities or developmental speech/language impairment, or taking psychotropic medication within the last two years. All participants signed a written consent before taking part in the study and were reimbursed by course credit or gift vouchers upon completion of the study. The study was approved by QUT Human Research Ethics Committee (LR 2023-5352-15049).

### 2.2 Stimuli

We selected six categories of animals (birds, felines, dogs, fish, invertebrates, and rodents) and five categories of man-made objects (furniture, gardening tools, kitchen utensils, musical instruments, and vehicles). These categories were defined as being either the reference category (birds), the low distance (LD) categories (as compared to the reference, i.e., other basic categories of animals) and the high distance (HD) categories (categories of man-made objects). For each sub-category, we selected 10 exemplar words (e.g., cuckoo, pelican, owl, swan, raven, eagle, goose, dove, duck and canary for the birds category; see **Supplementary Material** for all word stimuli) so that words in each of the stimulus sets (reference, LD and HD) were matched on length (number of letters), frequency of orthographic form, number and frequency of orthographic neighbours, bigram and trigram frequency as retrieved from the MCWord database (Medler & Binder, 2005; https://www.neuro.mcw.edu/mcword/) (**Table 1**). The matching procedure between stimulus sets was performed using the Match software (van Casteren & Davis, 2007). To ensure the matching procedure was successful, we computed independent-samples two-tailed t-tests comparing reference stimuli to LD or HD stimuli, and LD to HD stimuli on these six psycholinguistic variables. All p-values ranged between 0.15 and 0.91, indicating no significant differences between sets.

**Table 1.** Psycholinguistic characteristics of the word exemplars in each category. Mean (standard deviation).

|  | **Reference category**<br>**(n = 10)** | **LD category**<br>**(n = 50)** | **HD category (n**<br>**= 50)** |
| --- | --- | --- | --- |
| **Length** | 4.9 (1.2) | 5.7 (1.6) | 5.6 (1.4) |
| **Freq. of orthographic form** | 4.2 (2.9) | 3.9 (5.5) | 6.4 (8.9) |
| **N. of orthographic neighbours** | 5.3 (5.8) | 3.6 (6.2) | 4.1 (4.6) |
| **Freq. of orthographic**<br>**neighbours** | 35.6 (82.6) | 98.6 (607.7) | 29.9 (77.4) |
| **Bigram freq.** | 1158.6 (884.3) | 981.8 (926.8) | 826.1 (710.0) |
| **Trigram freq.** | 140.3 (160.9) | 127.7 (188.9) | 133.3 (179.5) |

Note: Freq. = frequency; N. = number.

For the reference category (birds) and for each exemplar we created five picture stimuli from natural colour photographs taken from the internet and resized to 300 x 300 pixels, and five word stimuli by writing the exemplar name in various fonts (Eras Medium ITC, MV Boli, Tahoma, Bradley Hand ITC, and Ink Free) and colours (yellow, black, blue, green, and purple) on a light grey background, for a total of 50 picture and 50 word stimuli. For all the other categories, we created four stimuli of each type (picture or word) following the same procedure as the reference category, for a total of 200 picture and 200 word stimuli per category type. Note that unlike some recent previous FPVS studies using pictures (e.g., Peykarjou et al., 2024b, 2024a; Stothart et al., 2017), we chose to include natural photographs of items, not cropped from their background, to reduce the contribution of low-level contour effects.

We also created control stimuli for each picture and word stimuli. For pictures, each original stimulus was distorted using diffeomorphic transformation until reaching level 25 of warping at which images from categories such as birds, animals or different kinds of objects have been found to become unrecognizable by human observers (see picture stimuli in the Pic Ctrl HD and Pic Ctrl LD conditions in **Figure 1**; Stojanoski & Cusack, 2014, https://github.com/rhodricusack/diffeomorph). For words, each original stimulus was vertically inverted (e.g., Word Ctrl LD in **Figure 1**) based on a recent study showing that vertically inverted words (as compared to 180° rotated or upright words) most significantly impact word recognition processes (Sussman et al., 2018).

**Figure 1.**
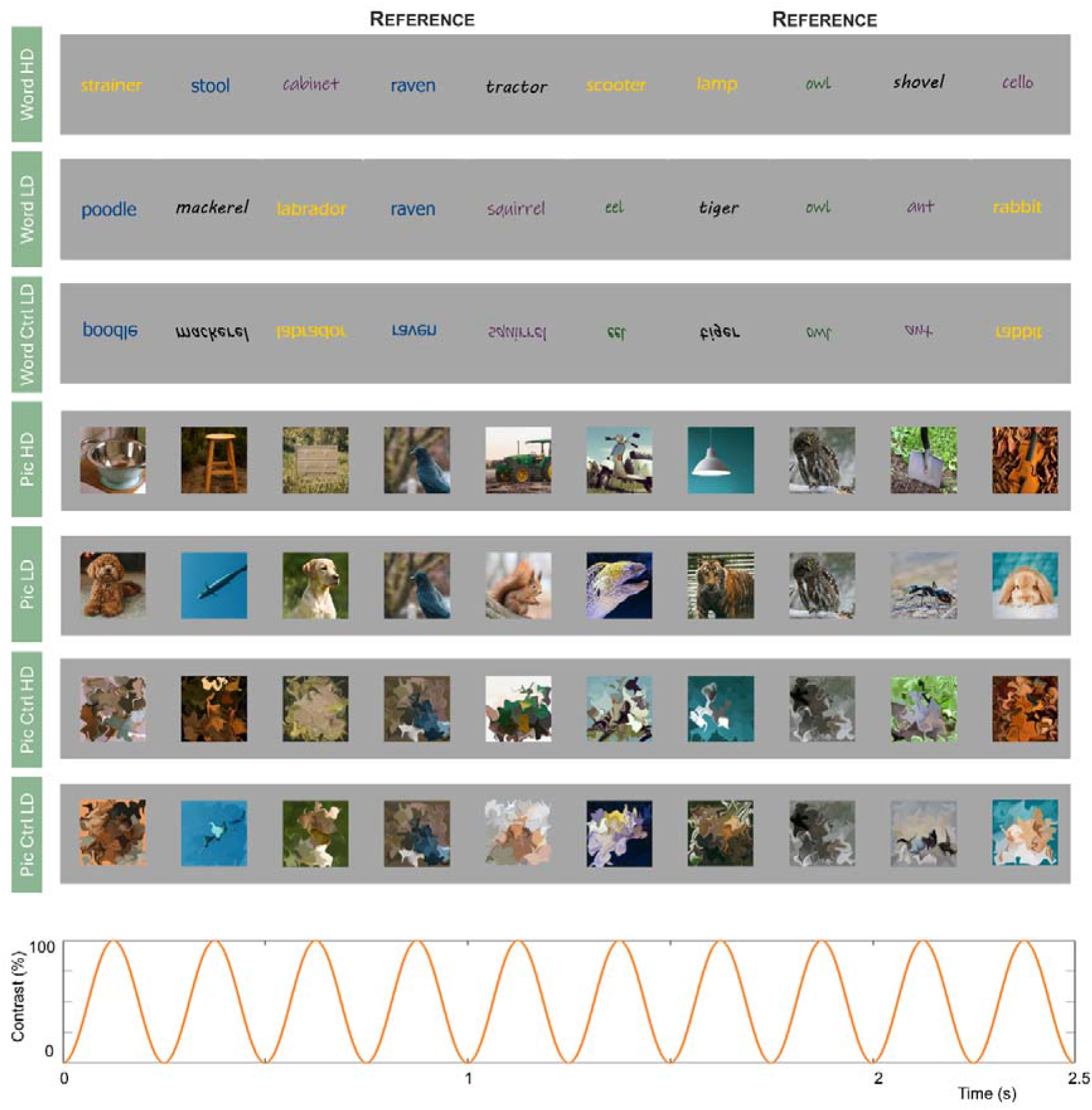
Schematic representation of the fast periodic visual stimulation sequences in the picture and word conditions. Pictures or words from the LD or HD stimulus categories were presented at a frequency of 4 Hz (i.e., 4 stimuli per second) using sinusoidal contrast modulation, and a picture or word from the reference category (i.e., birds) was inserted every 4 items (i.e., at 1 Hz). The size of each stimulus varied randomly from 80 to 120% of their original size at each cycle (not represented here for visualization purposes). An example of control conditions is provided. Note that the pictures used in this figure were not those used in the original paradigm for copyright reasons. Ctrl = Control conditions; HD = High Distance conditions; LD = Low Distance conditions; Pic = Picture conditions.

### 2.3 Procedure

#### 2.3.1 Scalp EEG recordings

Each participant was tested individually in a dimly lit room, seated at 80 cm from the computer screen while their electrical brain activity was recorded through a 61-channels custom equidistant hexagonal ANT Neuro EEG system. EOG was recorded by adding one electrode below the left eye and one electrode at its outer canthus. Scalp EEG signal was sampled at 512 Hz and electrodes were referenced to a central electrode (5Z) with the ground electrode taped to the left collar bone of the participant.

#### 2.3.2 Fast periodic visual stimulation

The experiment was run in a custom software running on Java and included 8 conditions: two conditions with picture stimuli (Pic LD and Pic HD), two conditions with control picture stimuli (Pic Ctrl LD and Pic Ctrl HD), two conditions with word stimuli (Word LD and Word HD) and two conditions with control word stimuli (Word Ctrl LD and Word Ctrl HD). For each stimulus type, one condition contrasted the LD category items with the reference category items while the other contrasted the HD category items with the reference ones. Parallel to previous neuropsychological explorations of semantic distance (T. T. Rogers et al., 2015), the same reference items were used across HD and LD conditions to ensure that these item characteristics were perfectly matched across conditions. For each condition, stimuli from the LD or HD categories were presented at a frequency rate of 4 Hz (i.e., 250 ms per stimulus) through sinusoidal contrast modulation, with the contrast of each stimulus gradually increasing from 0% to reach 100% and then decreasing to 0%. Every 4 stimuli, a stimulus belonging to the reference category appeared, i.e., at 1 Hz (**Figure 1**). To minimize the influence of low-level features, the stimulus size varied randomly between 80 and 120% of the original stimulus size in steps of 10%. At an 80-cm distance from screen, the visual angle of words thus ranged from 2.5 to 10° in width and from 1.1 to 1.9° in height while the visual angle of pictures varied from 4.7 to 7.1° in width and height. LD and HD stimuli were pseudo-randomly selected within each category to avoid consecutive presentation of the same exemplar (e.g., no two “axe” items after each other). Reference stimuli were randomly selected within their corresponding stimulus folder.

Stimuli were presented in sequences of 70 seconds, with an additional 2 seconds of contrast fade-in at the start of the sequence and 2 seconds of fade-out at the end. In line with previous studies using a similar paradigm (e.g., Peykarjou et al., 2024b; Rossion et al., 2015; Volfart et al., 2021), four sequences of each condition were presented, for a total here of 32 sequences per participant. The order of sequences was pseudo-randomized in two blocks of 16 sequences for every participant, with a break in-between.

The participant’s task was to focus their attention on the stimuli presented while also attending to two vertical fixation bars located on each side of the stimulus. The fixation bars randomly changed colour 8 times during each sequence, changing from black to white for 500 ms either one at a time or both at the same time. The participant was asked to press the space bar when the two fixation bars changed colour at the same time but to refrain from responding if only one fixation bar changed colour. This task recruiting more deployed spatial attention than a central fixation task (as used in previous FPVS studies, e.g., Rossion et al., 2015; Stothart et al., 2017; Volfart et al., 2021) was recently shown to elicit stronger electrophysiological responses in a FPVS paradigm investigating word-selective responses (Lochy et al., 2024).

#### 2.3.3 Vocabulary task

As some of the word stimuli used in the FPVS experiment were of low frequency, we also conducted a 10-min vocabulary task after EEG recordings to assess the participants’ conceptual knowledge of each word. The idea behind this task was to be able to correlate its results with EEG responses, the hypothesis being that a participant with less knowledge of words presented during the FPVS task may exhibit smaller EEG amplitudes in word conditions.

The EEG cap was removed, and the participant was presented with a behavioural task programmed in PsychoPy (version 2022.2.5; https://www.psychopy.org/). Each of the 110 word stimuli used in the FPVS experiment (belonging to 11 different categories) and 12 filler words (i.e., not presented during the FPVS experiment; 3 words from 4 different categories, i.e., clothing items, fruits, weapons and reptiles) was presented individually on the screen in random order. For each word, the participant was asked to: 1) indicate whether they know the word or not (by clicking “yes” or “no” on the screen), 2) estimate how confident they are about knowing this word’s meaning (by selecting a rating on a 5-point slider scale from 1, “not very confident”, to 5, “very confident”), and 3) indicate to which semantic category they think the word belongs to (by clicking on a category name among 15 presented simultaneously on the screen). If the participant clicked “no” at the first question, they were directly presented with the next word without going through the two other questions. Questions remained on the screen for as long as the participant needed although they were asked to respond as quickly as possible.

Based on the behavioural responses at the vocabulary task, we calculated the percent of words rated as known, the mean confidence rating and the categorisation accuracy (the last measures being calculated only on experimental words rated as known).

### 2.4 EEG analysis

EEG analysis was conducted in MATLAB 2018b (The MathWorks, Inc.) and the Letswave 6 toolbox (https://github.com/NOCIONS/letswave6).

#### 2.4.1 Preprocessing

Raw EEG datafiles were imported and continuous blocks of recordings were aligned to the first block. After Butterworth bandpass filtering between 0.1 and 100 Hz and notch filtering at 50 Hz (0.5 Hz width), the EEG signal was downsampled to 256 Hz to reduce file size and processing time. The datafile was then segmented according to event triggers of each condition, including 2 seconds before and after each sequence.

An independent component analysis (ICA) was computed using the runica algorithm and a single component capturing eye blink artefacts was removed for all participants. Residual noisy electrodes were replaced using linear interpolation (mean: 1.08 ± 1.2). All electrodes were re-referenced to the common average and datafiles were segmented for the second time using an integer number of 1 Hz cycles, excluding the fade-in and fade-out portions of each sequence (resulting in 17,682 bins, i.e., 69.0703 sec). For each participant, this led to 4 epochs per condition that were then averaged before being transformed to the frequency domain.

#### 2.4.2 Frequency-domain analyses

A Fast Fourier Transform (FFT) was applied to these averaged epochs and amplitude spectra were extracted for each individual. We computed three types of baseline correction transforms to compare each frequency bin to its 48 neighbouring bins (24 bins on each side, excluding the immediately adjacent bin): 1) EEG signal in the neighbouring bins was subtracted from each frequency bin to quantify the electrophysiological response in microvolts across harmonics; 2) each frequency bin was divided by the EEG signal in the neighbouring bins to estimate the signal-to-noise ratio (SNR); and 3) z-scores were computed as the difference between the amplitude at each frequency bin and the mean amplitude in its neighbouring bins, divided by the standard deviation of the neighbouring bins.

To quantify the response distributed across harmonics of the frequency of interest (1 Hz), we computed the sum of harmonics by segmenting each FFT epoch into successive chunks of 51 bins centred around 1 Hz and harmonics up until 7 Hz. The choice of harmonics to consider for the sum was made based on the consecutive harmonics showing a significant response when all conditions and participants were grand-averaged. These 6 epochs (centred around 1, 2, 3, 5, 6 and 7 Hz) were summed together, excluding the epoch centred around 4 Hz, representing the base frequency, and baseline correction transforms were applied. Individual results were extracted for each electrode and the data were grand-averaged to get scalp topographies of activation.

We also computed the sum of harmonics of the base frequency (4 Hz) based on the same principle as described above, i.e., looking at significant consecutive harmonics over the grand-average of conditions and participants. In this case, we summed successive chunks of 51 bins centred around 4 Hz and harmonics up until 28 Hz (7^th^ harmonic of the base frequency) and computed baseline correction transforms for quantification of the responses.

#### 2.4.3 Definition of regions-of-interest

We defined three regions of interest (ROI) for subsequent analyses. Two were in the bilateral occipito-temporal (OT) ROIs and were defined based on regions highlighted in a previous FPVS study investigating semantic word categorization responses (Volfart et al., 2021) and FPVS studies using lexical stimuli (e.g., Lochy et al., 2024). Due to the use of equidistant EEG caps (as opposed to caps following the 10/20 electrode system in previous studies), we selected 5 occipito-temporal electrodes on each side (LOT and ROT) that approximately overlapped with the OT ROIs described in the abovementioned studies. A third ROI was defined over the centro-parietal region (6 electrodes) to overlap with the locus of activity described in studies investigating semantic effects in scalp EEG, usually reflected by the N400 component (e.g., Kutas & Federmeier, 2011; see also David et al., 2025; Meyer et al., 2024).

### 2.5 Statistical analysis

Results at the behavioural orthogonal fixation task were analysed in terms of percent corrected accuracy (correcting for false alarms, i.e., button press to one bar or to nothing, using the formula: (N Hits / (N Hits + N False Alarms)) * ((N Hits / N Colour Change Two Bars) * 100)) and correct response times in milliseconds (RT). The behavioural results at the fixation bar task in each condition were compared in SPSS (IBM SPSS Statistics, version 28.0.1.0) using repeated measures ANOVAs.

The amplitudes of electrophysiological responses in each condition at the frequencies of interest were compared in SPSS using repeated measures ANOVAs. Where significant main effects or interactions were found, post hoc pairwise comparisons were conducted using paired t-tests. A p-value under 0.05 was used as statistical threshold for significance. Although some conditions deviated from normality based on Shapiro-Wilk tests, repeated measures ANOVAs were retained because they have been shown to be generally robust to violations of normality when the sphericity assumption is met (Blanca et al., 2023). Mauchly’s tests indicated that the assumption of sphericity was met for the ROI factor and all interactions involving ROI (all p-values > .05). Since all other factors in the present study had only two levels, the sphericity assumption was automatically satisfied for these factors. To further verify that our conclusions were not dependent on parametric assumptions and assess the robustness of the findings, additional Wilcoxon signed-rank tests were conducted for the principal HD-LD comparisons in each ROI.

The mean percent of words rated as known, the mean confidence rating and the mean categorisation accuracy of each participant in the vocabulary task were correlated with their EEG responses at the sum of harmonics of the reference frequency in each experimental condition using Pearson correlation coefficients. Correlations were corrected for multiple comparisons using FDR correction within conditions and ROI (Benjamini & Hochberg, 1995).

## 3 Results

### 3.1 Behavioural responses

#### 3.1.1 Fixation bars task

Response times below 150 and above 2000ms were considered as inaccurate and excluded from RT analysis. Mean percent correct accuracy and correct RT in each condition are displayed in **Table 2**.

**Table 2.** Behavioural results at the orthogonal fixation bars task. Mean ± SD.

|  | Mean corrected accuracy (%) | Mean RT (ms) |
| --- | --- | --- |
| Pic HD | 93.5 ± 6.5 | 528 ± 55 |
| Pic LD | 91.7 ± 7.5 | 513 ± 53 |
| Pic Ctrl HD | 94.4 ± 7.3 | 512 ± 59 |
| Pic Ctrl LD | 93.9 ± 6.4 | 504 ± 41 |
| Word HD | 94.0 ± 7.3 | 502 ± 54 |
| Word LD | 92.5 ± 8.0 | 498 ± 54 |
| Word Ctrl HD | 94.2 ± 6.8 | 510 ± 43 |
| Word Ctrl LD | 94.6 ± 6.9 | 509 ± 52 |

Note: Ctrl = Control conditions; HD = High Distance conditions; LD = Low Distance conditions; Pic = Picture conditions; RT = response time.

A repeated measures ANOVA with factors Item Type (Pictures vs. Words), Condition Type (Control vs. Experimental) and Distance (HD vs. LD) was computed to compare corrected accuracy across conditions. It showed only a trend towards significance for the main effect of Condition Type (F(1,23) = 3.645, p = 0.069). Other main effects did not reach significance (Item Type: F(1,23) = 0.271, p = 0.607; Distance: F(1,23) = 1.137, p = 0.297). None of the interaction effects reached significance (F range: 0.020 to 0.626; p range: 0.437 to 0.889).

A similar ANOVA was computed on correct RT and showed a significant effect of Item Type (F(1,23) = 4.932, p = 0.036), indicating longer response times in picture than word conditions. Other main effects were not significant (Condition Type: F(1,23) = 0.255, p = 0.618; Distance: F(1,23) = 2.108, p = 0.160). None of the interaction effect was significant (F range: 0.032 to 1.282, p range: 0.269 to 0.860) except for the interaction effect Item Type * Condition Type (F(1,23) = 15.730, p < 0.001), suggesting that RT were longer in control than experimental conditions for words but longer for experimental than control conditions for pictures.

#### 3.1.2 Vocabulary task

The mean percent of words rated as known across all participants was 95.2 ± 3.8 %, suggesting that a high proportion of words were known to all participants. The mean confidence rating (out of 5, “very confident”) was 4.6 ± 0.2, indicating that participants were highly confident in knowing what the words meant. The mean categorisation accuracy was 96.2 ± 2.4 %, suggesting that the participants were able to select the correct semantic category for the majority of words. The results indicate that participants were highly familiar with most of the word stimuli presented during the FPVS experiment.

### 3.2 EEG responses

#### 3.2.1 Response at the reference category frequency

**Whole scalp analyses.** Electrophysiological responses at the reference category frequency (i.e., 1 Hz) were significant up to the 7^th^ harmonic (7 Hz) when considering the average of conditions and participants. The topographical distribution of EEG responses at the sum of these harmonics highlighted similar topographies across experimental conditions, with clear bilateral occipito-temporal foci of activation for both pictures and words (**Figure 2**).

**Figure 2.**
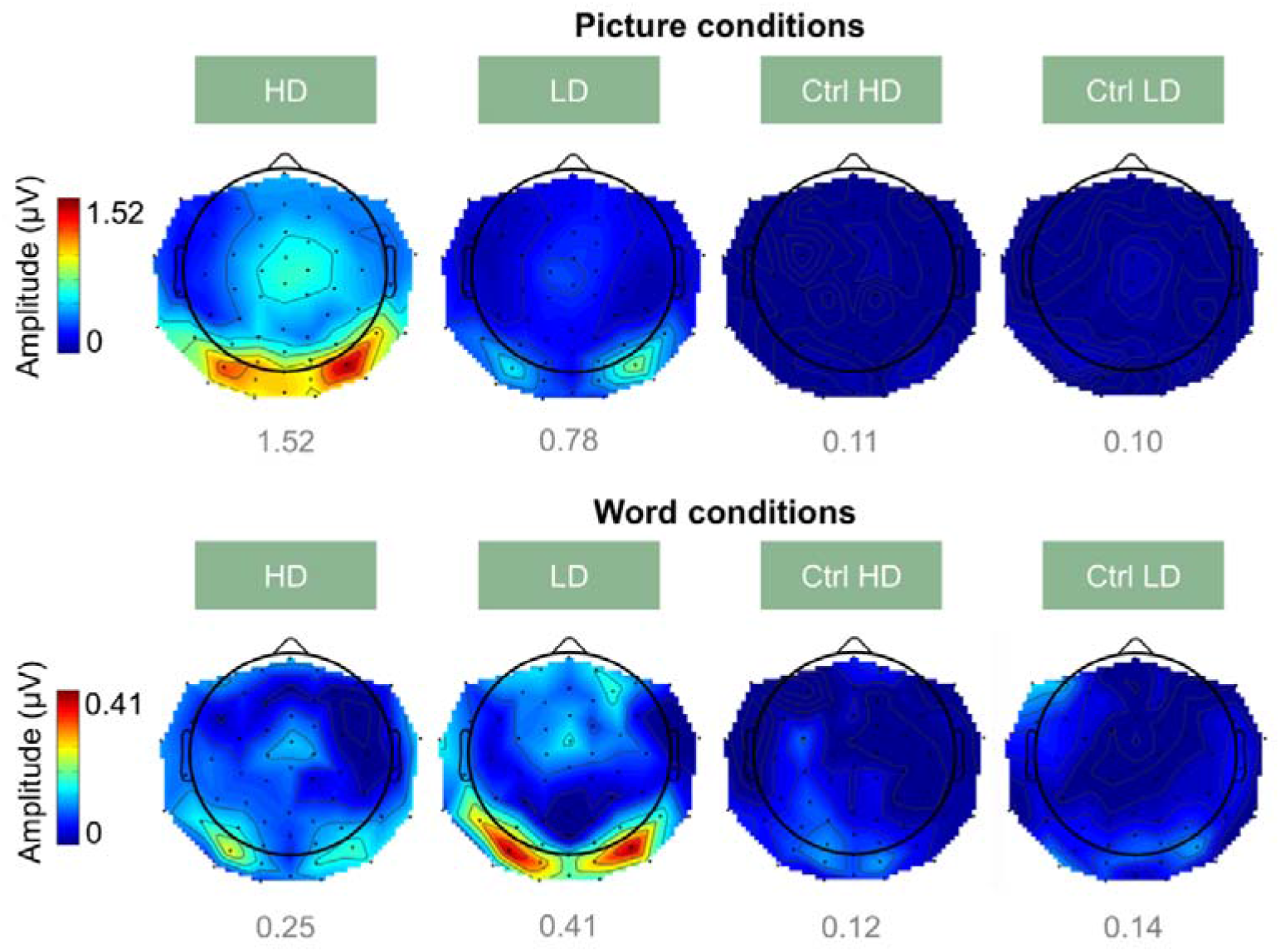
Topographical distribution of baseline-corrected amplitudes at the sum of harmonics of the reference category frequency (grand-averaged data, n = 24 participants). The colour scale is adjusted to the maximal amplitude across picture and word conditions separately. The maximal amplitude in a given condition is shown in grey below each map.

We compared the whole scalp response at the sum of harmonics across conditions using a repeated measures ANOVA with factors Item Type (Pictures vs. Words), Condition Type (Control vs. Experimental) and Distance (HD vs. LD). This analysis showed a main effect of Item Type (F(1,23) = 40.994, p < 0.001), Condition Type (F(1,23) = 101.018, p < 0.001) and Distance (F(1,23) = 12.222, p = 0.002). All interaction effects were significant (Item Type * Condition Type: F(1,23) = 119.521, p < 0.001; Item Type * Distance: F(1,23) = 23.987, p < 0.001; Condition Type * Distance: F(1,23) = 28.093, p < 0.001; Item Type * Condition Type * Distance: F(1,23) = 38.490, p < 0.001). These effects indicated marked differences between all conditions that was further analysed at the level of ROI.

**ROI analyses.** Frequency spectra in each ROI and condition are shown in **Figure 3**. Mean amplitudes in each ROI and condition are displayed in **Figure 4**.

**Figure 3.**
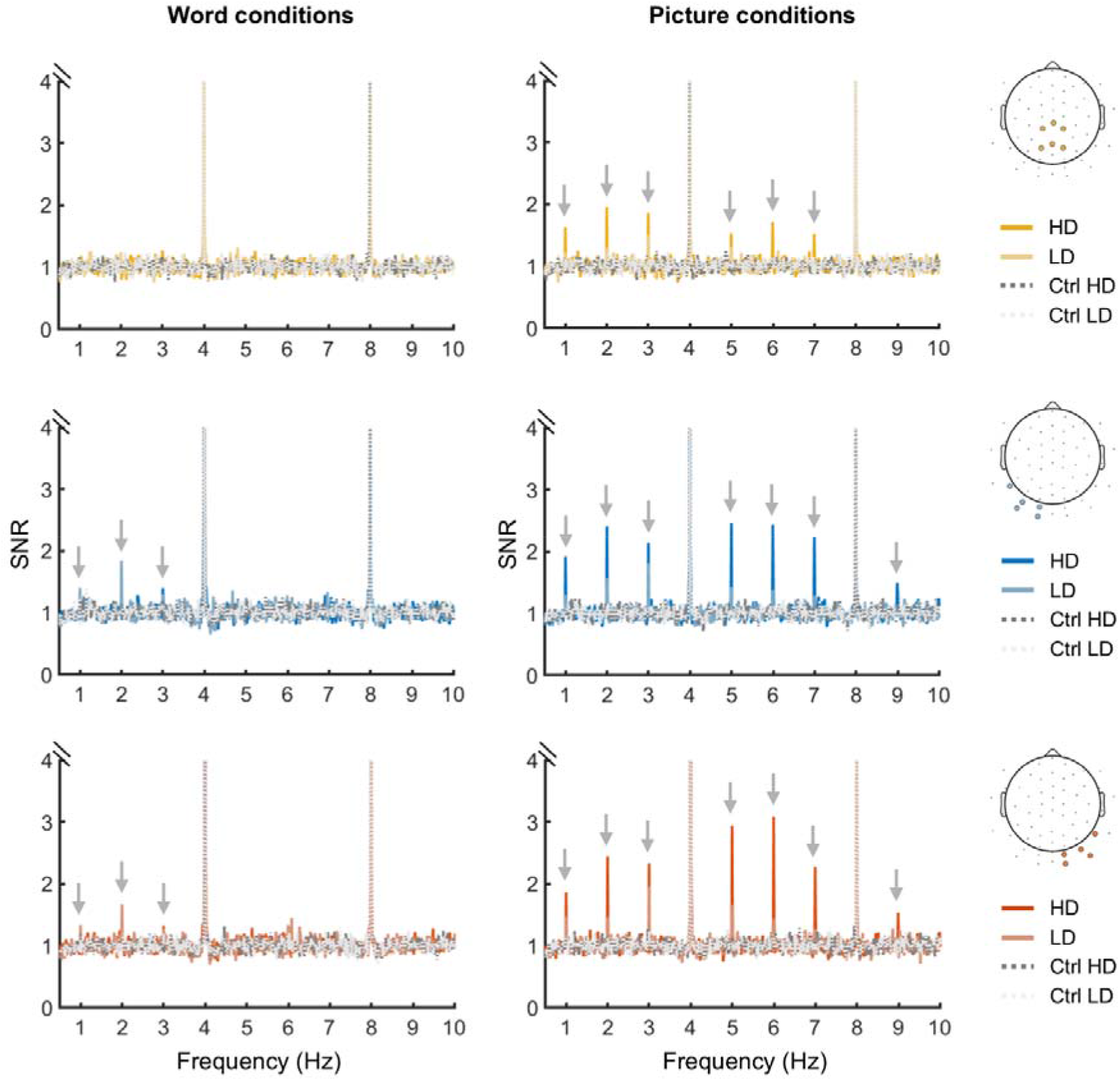
Frequency spectra of the signal-to-noise ratio in each ROI for word and picture conditions. Channels included in each ROI are shown on the right with coloured circles on the head template and each spectrum displays the average SNR in each ROI. Grey arrows point to responses at the reference category frequency (i.e., 1 Hz) and its harmonics. The y-axis is cut at 4 for visualization purposes as the base frequency (i.e., 4 Hz and its harmonic 8 Hz) have much higher SNR than the reference category frequency.

**Figure 4.**
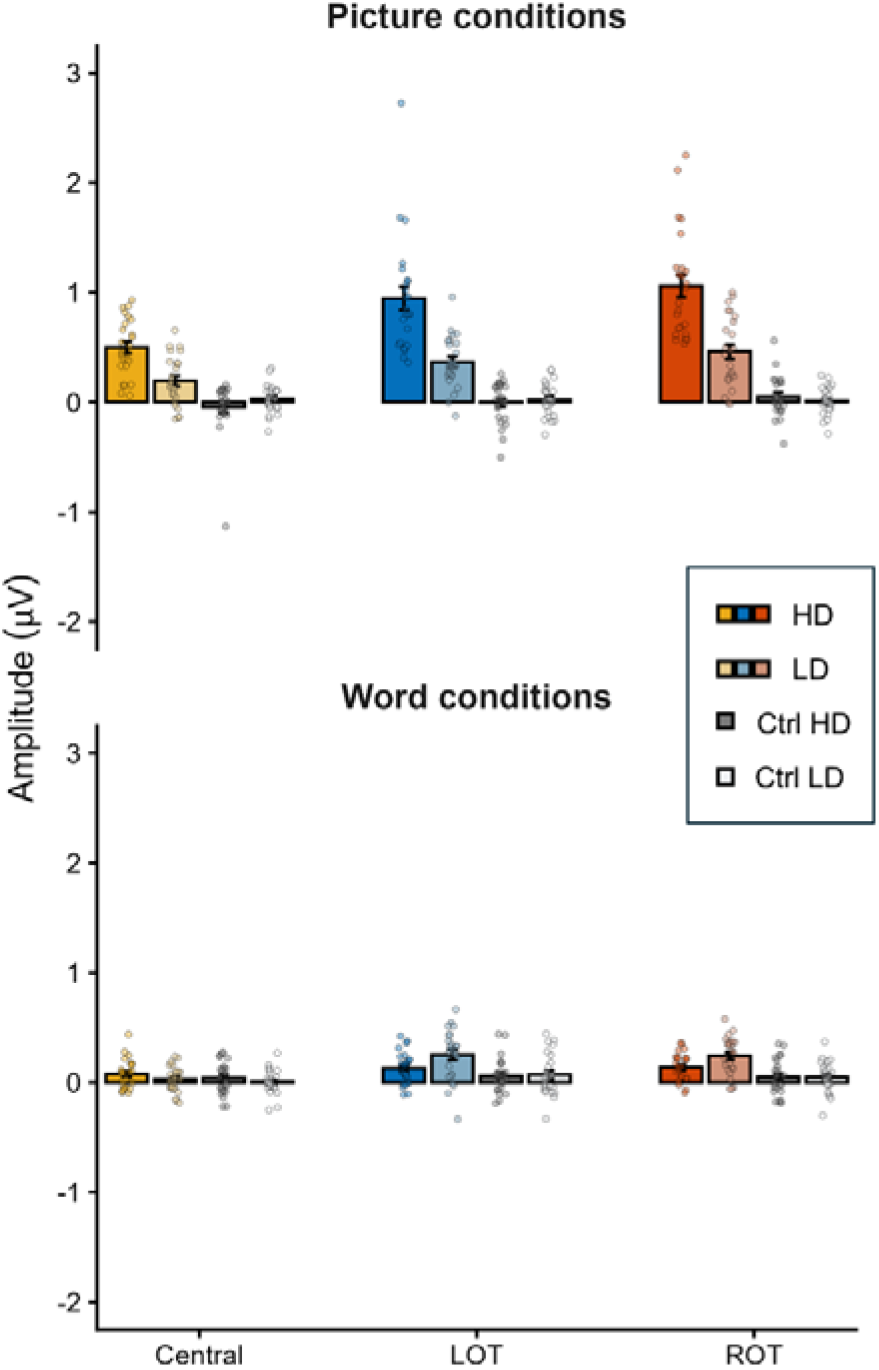
Baseline-corrected amplitudes at the sum of harmonics of the reference category frequency in each ROI for all picture and word conditions. Bars represent the mean amplitude across participants, coloured dots represent individual participant amplitudes, and error bars represent ±1 standard error from the mean.

Z-scores were computed as described in **Section 2.4.2** on the average of participants and channels within each ROI and for each condition to assess significance of the response at the reference frequency. Over the LOT and ROT ROI, all experimental but none of the control conditions showed a significant response at the reference frequency. Over the central ROI, only responses in the HD and LD picture conditions and HD word conditions were significant (z > 1.65).

We conducted a 2 (Item Type) x 2 (Condition Type) x 2 (Distance) x 3 (ROI) repeated measures ANOVA on the sum of harmonics at the reference category frequency. All main effects were significant (Item Type: F(1,23) = 45.862, p < 0.001; Condition Type: F(1,23) = 129.019, p < 0.001; Distance: F(1,23) = 15.026, p < 0.001; ROI: F(2,46) = 34.636, p < 0.001). All first-order interaction effects were significant (Item Type * Condition Type: F(1,23) = 129.660, p < 0.001; Item Type * Distance: F(1,23) = 20.675, p < 0.001; Condition Type * Distance: F(1,23) = 20.840, p < 0.001; Item Type * ROI: F(2,46) = 9.843, p < 0.001; Condition Type * ROI: F(2,46) = 32.358, p < 0.001) except for the interaction effect Distance * ROI (F(2,46) = 0.945, p = 0.396). All second-order interaction effects were significant (Item Type * Condition Type * Distance: F(1,23) = 33.491, p < 0.001; Item Type * Condition Type * ROI: F(2,46) = 9.303, p < 0.001; Item Type * Distance * ROI: F(2,46) = 10.047, p < 0.001) except for the Condition Type * Distance * ROI interaction effect (F(2,46) = 0.351, p = 0.706). The third-order interaction effect did not reach significance (Item Type * Condition Type * Distance * ROI: F(2,46) = 2.998, p = 0.060). These effects indicated strong differences between pictures and words, control and experimental conditions, HD and LD, and the three ROIs.

To better investigate the effects of interest and considering the strong difference in amplitude between pictures and words, we computed repeated measures ANOVAs separately on picture and word experimental conditions with factors Distance (LD vs. HD) and ROI (central, LOT and ROT). For pictures, both main effects were significant (Distance: F(1,23) = 30.867, p < 0.001; ROI: F(2,46) = 35.314, p < 0.001), as well as the interaction effect (F(2,46) = 4.635, p = 0.015). Pairwise comparisons between ROI highlighted lower amplitudes in the central ROI than in the LOT and ROT (p < 0.001 for both comparisons), without difference between the two OT ROIs (p =0.120). Post-hoc paired t-tests assessing the effect of distance on each ROI separately showed a significant effect in all three ROIs (central: t(23) = 4.420, p < 0.001; LOT: t(23) = 4.885, p < 0.001; ROT: t(23) = 4.665, p < 0.001). For words, the main effect of ROI was significant (F(2,46) = 17.079, p < 0.001) while the main effect of Distance was not (F(1,23) = 3.489, p = 0.075). The interaction effect was significant (F(2,46) = 6.770, p = 0.003). Pairwise comparisons between ROI highlighted lower amplitudes in the central ROI than in the LOT and ROT (p < 0.001 for both comparisons), without difference between the two OT ROIs (p = 1). Post-hoc paired t-tests assessing the effect of distance on each ROI separately showed a significant effect over the LOT and ROT (LOT: t(23) = −2.649, p = 0.014; ROT: t(23) = −2.270, p = 0.033) but not over the central ROI (t(23) = 1.244, p = 0.226). These results indicated a significant distance effect in all ROI for picture conditions, with higher amplitudes in the HD than LD condition, while the distance effect was only significant for OT ROIs in word conditions, with higher amplitudes in the LD than HD condition (opposite direction to picture results).

To assess the robustness of the findings to departures from normality, HD-LD comparisons were additionally examined using Wilcoxon signed-rank tests. For picture stimuli, the HD condition elicited significantly larger responses than the LD condition in all three ROIs (central ROI: z = −3.37, p < .001; LOT: z = −3.89, p < .001; ROT: z = −3.91, p < .001). For word stimuli, responses were significantly larger in the LD than HD condition in the LOT (z = −2.54, p = .011) and ROT (z = −2.09, p = .037), but the comparison did not reach significance in the central ROI (z = −0.83, p = .407). These non-parametric analyses therefore confirmed the overall pattern of results obtained using parametric analyses.

Visual inspection of the individual data revealed two observations exceeding ±2.5 SD from the condition mean (from two different participants). Repeating the analyses after excluding these participants yielded the same overall pattern of results at both whole-scalp and ROI levels. The only difference was that the Distance * ROI interaction in the picture-only ROI analysis decreased from significance (*p* = .015) to trend level (*p* = .051).

#### 3.2.2 General stimulation response at 4 Hz

Electrophysiological responses at 4 Hz were significant up to the 7^th^ harmonic (28 Hz) when considering the average of conditions and participants. The sum of these harmonics (4 to 28 Hz) revealed that electrophysiological activity was maximal over bilateral occipito-temporal regions in all conditions (see **Supplementary Figure S1A**).

A repeated measures ANOVA with factors Item Type (Pictures vs. Words), Condition Type (Control vs. Experimental) and Distance (HD vs. LD) showed that the mean amplitudes across the whole scalp did vary slightly across conditions (see **Figure S1B**). There was a main effect of Condition Type (F(1,23) = 11.788, p = 0.002), with smaller amplitudes in the control than experimental conditions. There was also a main effect of Distance (F(1,23) = 6.931, p = 0.015), indicating smaller amplitudes in HD than LD conditions. The main effect of Item Type was not significant (F(1,23) = 0.368, p = 0.550). None of the interaction effects were significant (Item Type * Condition Type: F(1,23) = 0.247, p = 0.624; Item Type * Distance: F(1,23) = 1.293, p = 0.267; Condition Type * Distance: F(1,23) = 0.325, p = 0.574; Item Type * Condition Type * Distance: F(1,23) = 1.962, p = 0.175). These results suggested comparable amplitudes at the general stimulation frequency between picture and word conditions. However, they revealed lower amplitudes for control than experimental conditions and for HD than LD conditions.

#### 3.2.3 Correlations with vocabulary task responses

Across experimental conditions and ROI, we only observed significant correlations between EEG amplitudes in the Pic HD condition in the central ROI and the mean percent of words rated as known and the mean confidence rating (r = −0.52, FDR-corrected p = 0.036, and r = −0.48, FDR-corrected p = 0.036, respectively). None of the other correlation coefficients between vocabulary measures and EEG measures reached significance.

## 4 Discussion

The present study investigated whether semantic distance between concepts modulates electrophysiological responses to pictures and words using a FPVS paradigm. Across conditions, we observed robust EEG responses at the reference category frequency (1 Hz), indicating automatic and implicit discrimination of bird exemplars from other categories, which is consistent with previous FPVS-EEG studies contrasting object categories through written words (e.g., Volfart et al., 2021) or pictures (e.g., with superordinate, living/non-living categories, e.g., Peykarjou et al., 2024b, 2024a; but also more specific discriminations, e.g., faces, body parts and houses among various object types in Jacques et al., 2016; animals vs. fruit/vegetables or birds vs. mammals in Stothart et al., 2017). Critically, semantic distance effects emerged for picture stimuli, with significantly larger amplitudes in the high distance (HD) condition than in the low distance (LD) condition across all three regions of interest (LOT, ROT and central). In contrast, word stimuli elicited weaker overall responses, and the semantic distance effect was less consistent: a small reverse effect was observed in the occipito-temporal ROIs.

### 4.1 Picture stimuli lead to strong EEG discrimination responses and robust semantic distance effects

Our first research question asked whether exemplars from the same category would be automatically and implicitly discriminated from exemplars of other categories, regardless of the semantic distance. Our results clearly support such discrimination for picture stimuli: reference bird exemplars were robustly discriminated from base stimuli in both HD and LD conditions, with strong EEG responses at 1 Hz and harmonics. These results are concordant with previous FPVS studies using pictures to assess object discrimination at different levels (see e.g., Jacques et al., 2016; Peykarjou et al., 2024b, 2024a; Stothart et al., 2017).

Turning to our second research question asking whether we would observe a semantic distance effect in EEG discrimination responses, our findings are also in support of our hypothesis, showing a strong semantic distance effect for pictures, as manifested by significantly larger amplitudes in the HD condition than in the LD condition across all occipito-temporal and central ROIs. These findings of a strong semantic distance effect for pictures align well with the hub-and-spoke model (and other feature-based or vector-based models of semantic representations) but less so with strictly categorical accounts that posit discrete category boundaries and would predict category discrimination but not graded effects of semantic similarity (see Frisby et al., 2023 for a review). In particular, the hub-and-spoke model predicts that broader semantic distinctions (e.g., at the superordinate level, like living vs. non-living) emerge earlier during activation than more specific distinctions (e.g., dog vs. bird) and are also more robust under degraded and time-constrained conditions (McClelland & Rogers, 2003; T. T. Rogers et al., 2004; T. T. Rogers & Patterson, 2007; see also Cox et al., 2024; Lambon Ralph et al., 2017; T. T. Rogers et al., 2021). The predictions regarding performance under degraded conditions are notably informed by SD patient data which have shown relatively preserved performance for performing tasks at the superordinate level compared to basic or subordinate levels despite severe semantic impairment (e.g., T. T. Rogers et al., 2004, 2015; T. T. Rogers & Patterson, 2007). Conversely, an absence of semantic distance effects, that is, comparable responses for the HD and LD conditions, would have been difficult to reconcile with the graded similarity structure assumed by the hub-and-spoke framework and related similarity-based accounts.

In an attempt to better understand the discrepancy between this superordinate-over-basic advantage observed in SD patients and the basic-level advantage in healthy participants (e.g., Rosch et al., 1976), Rogers & Patterson (2007) also investigated performance under time constraints. In a picture category verification study imposing response deadlines to healthy participants, they found a decline in accuracy for basic and subordinate levels but not for the superordinate levels with increasing time constraints, replicating the pattern of performance observed in SD patients. More support for the robustness of superordinate distinctions come from behavioural studies using a rapid visual categorization task (stimulus presentation ∼20-30ms) that have demonstrated better performance at superordinate than basic or subordinate levels (e.g., Macé et al., 2009; Praß et al., 2013; see also Fabre-Thorpe, 2011). These previous studies are in line with our findings of greater EEG responses for superordinate compared to basic contrasts in the healthy population under the temporal constraints of the FPVS paradigm, where each stimulus was presented for only ∼250 ms before being masked by the following, allowing only limited time for semantic processing to unfold. Our results thus provide additional evidence in favour of a similarity-based semantic structure as postulated by the hub-and-spoke model, indicating that the time given to the system to process a stimulus influences neural activation pattern at a given point in time as the whole system dynamically settles towards the detailed semantic representation (e.g., Cox et al., 2024; T. T. Rogers et al., 2021).

### 4.2 Word stimuli show weaker EEG discrimination responses and less reliable semantic distance effects

Regarding our first research question and in contrast to results in picture conditions, word stimuli elicited significant but weaker EEG discrimination responses and a less consistent semantic distance effect, as initially predicted. A potential factor that may have contributed to the stronger responses in the LD condition is the greater semantic homogeneity of this set. When bird items were presented among other animal-related words (LD), it may be that the shared semantic context facilitated activation of all animal-related categories. In contrast, bird items were embedded in a more heterogeneous set of man-made objects in the HD condition, providing a less coherent semantic context priming that could have weakened discrimination responses. Such an interpretation could be related to the semantic priming literature, where activation of one concept is thought to spread to related concepts and sometimes facilitate semantic processing (e.g., Collins & Loftus, 1975; Levelt et al., 1999). In the context of semantic degradation as observed in semantic dementia, this literature has provided valuable insights into the organization of semantic memory. For instance, it is very well established that the patients’ global semantic degradation has its first and strongest effects on fine discriminations and distinctive semantic features (e.g., T. T. Rogers et al., 2015). Consistent results have also emerged from explorations of priming effects in this patient group. These investigations have shown that priming effects based on distinctive semantic features are diminished earlier than those based on shared semantic attributes or broader category relationships (e.g., Laisney et al., 2011; S. L. Rogers & Friedman, 2008). However, caution is warranted when comparing these priming findings to the present paradigm. Classical semantic priming effects have typically been studied using explicit behavioral tasks such as lexical decision, naming, or categorization (e.g., Bajo, 1988; Hantsch et al., 2012; Howard et al., 2006; Vigliocco et al., 2002), and have been shown to depend on the task used, with semantic context effects often reduced or absent in tasks that can be solved without conceptual processing (Belke, 2013; Wöhner et al., 2024). In contrast, the FPVS paradigm employed here involved rapid, continuous stimulus presentation and semantic activation without requiring any explicit semantic task or decisions. Therefore, while semantic homogeneity may have contributed to the larger responses observed in the LD word condition, the extent to which semantic priming mechanisms account for this effect remains unclear.

The pattern of results observed for the semantic distance effect is consistent with our second hypothesis and with predictions from the hub-and-spoke model which posits a difference in mapping between picture or word inputs and semantic representations (Lambon Ralph & Howard, 2000; T. T. Rogers et al., 2004; see also Caramazza et al., 1990 who similarly make the assumption of a “privileged access to meaning” for pictures but not words). Again, these predictions are based on SD patient data which usually highlight worse performance with word/naming than picture/comprehension tasks (e.g., Lambon Ralph et al., 2001, see also 2017; Patterson et al., 2007) and which have shown semantic distance effects for words collapsing rapidly with increasing impairment, while picture-based effects remain relatively stable with disease progression (Lambon Ralph & Howard, 2000; T. T. Rogers et al., 2004). Computational simulations that have implemented such difference in mapping to semantic representations by modelling picture input as patterns of activity across several units but modelling word input as either entirely arbitrary (Lambon Ralph & Howard, 2000) or as single units (T. T. Rogers et al., 2004) successfully replicated the pattern of performance observed in SD patients, with word comprehension deficits being disproportionately pronounced following damage to the semantic system.

Considering that a certain amount of time is needed for semantic representations to settle into their final, stable activation patterns (e.g., Cox et al., 2024; T. T. Rogers et al., 2021) and that the relationship between semantic representations and word inputs is largely arbitrary (e.g., Lambon Ralph & Howard, 2000; T. T. Rogers et al., 2004), it is possible that the rapid presentation rate used in our FPVS paradigm was more detrimental to word than picture processing by limiting the time available for semantic activation to unfold.

Moreover, as already mentioned above, the findings of Rogers and Patterson (2007) suggest that healthy participants under time pressure may behave similarly to SD patients, i.e., relying on poorly specified semantic representations, which have been demonstrated to disproportionately impair word more than picture processing (Lambon Ralph & Howard, 2000; T. T. Rogers et al., 2004). Our findings of weaker effects in word conditions suggest that our FPVS paradigm may have mimicked the conditions of semantic degradation, particularly for word stimuli, and highlights the limitations of FPVS for probing fine-grained semantic effects such as the semantic distance effect in written language, at least with a frequency of 4 Hz as used here. This possibility should be further explored in the future by slowing down stimulus presentation frequency and assessing the effect of this frequency change on picture and word conditions.

### 4.3 Contribution of visual features to semantic discrimination responses in picture conditions

While semantic effects were robust for pictures, it is important to acknowledge the potential contribution of visual features to EEG discrimination responses. Information about object identities or their taxonomic categories can be derived from visual properties (e.g., Eger et al., 2008; Gerlach et al., 2015; Iordan et al., 2015; Victoria et al., 2019), and spatial frequency availability has been shown to influence object grouping depending on the level (i.e., superordinate, basic or specific) at which a task must be performed (Ashtiani et al., 2017). More generally, semantic category membership and visual structure are strongly correlated in natural images: categories differing in animacy, taxonomic membership, or object function often exhibit systematic differences in image statistics, including spatial frequency distributions, orientation content, colour and background composition (Torralba & Oliva, 2003). Consequently, when using photographs of real-world objects, semantic and visual contributions are intricated. Although the present study included diffeomorphic control stimuli specifically chosen because they preserve perceptual image properties more faithfully than traditional scrambling procedures while disrupting recognizability (Stojanoski & Cusack, 2014), these controls cannot fully eliminate all category-related visual regularities. Accordingly, the semantic distance effect observed for pictures should not be interpreted as reflecting semantic processing in complete isolation from visual information. Rather, it likely reflects the processing of semantic distinctions as they occur in naturally structured visual object representations. This interpretation is consistent with recent accounts of semantic cognition, which view conceptual representations as intrinsically linked to surface level features (e.g., Dilkina & Lambon Ralph, 2013; Fernandino et al., 2022; T. T. Rogers et al., 2004; see also Lambon Ralph et al., 2017).

Importantly, the bilateral occipito-temporal and central ROIs implicated in our findings are consistent with regions known to support object recognition and semantic processing. Notably, the central region overlaps with scalp distributions traditionally associated with the N400 component, which has been linked to semantic access and integration in previous EEG studies (e.g., Gallagher et al., 2014; Kutas & Federmeier, 2011). This interpretation is further supported by recent evidence implicating the anterior temporal regions in N400 generation (Meyer et al., 2024), in line with proposals that these regions function as a semantic hub (Lambon Ralph, 2014; Lambon Ralph et al., 2017). This finding suggests that the observed responses likely reflect not only visual object discrimination but also semantic processes. Nevertheless, caution is warranted when relating our scalp-level EEG responses to the neural substrates implicated in semantic dementia because the posterior scalp distributions observed here cannot be directly linked to activity in the anterior temporal lobes. Future studies employing source localization of EEG/MEG measurements or intracranial recordings will be required to clarify this relationship.

### 4.4 FPVS as a promising tool for probing semantic architecture and clinical applications

Despite limitations for highlighting semantic distance effects in the word conditions, FPVS successfully elicited reference category frequency responses for both pictures and words, and HD and LD conditions, confirming its utility for probing implicit semantic discrimination processes. This approach complements previous FPVS studies (Stothart et al., 2017; Volfart et al., 2021) and offers a valuable tool for investigating semantic processing without requiring explicit tasks.

FPVS is particularly promising for use in clinical populations, such as patients with SD or Alzheimer’s disease, where fatigue, attentional and executive demands may hinder performance in more traditional, behavioural paradigms. A recent study using a similar semantic word discrimination paradigm have notably highlighted lower EEG response amplitudes at the frequency of category change in a sample of patients with Alzheimer’s disease as compared to age-matched controls, opening the way for the assessment of semantic categorization abilities and FPVS diagnosticity across various populations (David et al., 2025; see also Hermann et al., 2025; Stothart et al., 2021, 2025). To assess potential differences depending on the location of the brain lesions, future studies should for example explore how other brain-lesioned populations perform at different FPVS paradigms aimed to assess coarse or finer-grained, semantic or non-semantic functions.

### 4.5 Limitations

Several limitations should be acknowledged. First, our choice of image transformation to create control stimuli (diffeomorphic distortion vs. vertical inversion) may be questioned. While it was motivated by previous studies showing their efficiency in altering stimulus recognizability (i.e., the object’s meaning) but preserving important low-level perceptual features (Stojanoski & Cusack, 2014; Sussman et al., 2018, for pictures and words, respectively), the image transformation being different for pictures and words limits the comparability across stimulus types.

Second, while effort was made to match the word stimuli on several psycholinguistic variables across stimulus sets (reference, LD and HD), slight differences across sets could not be entirely eliminated. For instance, LD words are numerically less frequent and have fewer orthographic neighbours than the two other stimulus sets, but their neighbours are on average more frequent (although not significantly). Such factors could influence lexical activation and thus EEG discrimination responses, potentially amplifying contrasts if some LD words are less familiar and processed more like pseudowords. Future research should further examine how such differences on various psycholinguistic variables may influence EEG responses.

Finally, the rapid presentation rate in FPVS may have prevented semantic activation to reach its final activation pattern, affecting words more strongly due to reasons discussed above. Slowing the base rate (e.g., at 2.5 Hz, i.e., each stimulus would appear for 400ms) could allow deeper semantic processing and potentially reveal semantic distance effects in word conditions.

## 5 Conclusion

In conclusion, the present findings extend and complement previous FPVS studies suggesting that this objective electrophysiological approach can capture implicit semantic discrimination responses from both picture and word stimuli within a few minutes of recordings. We found robust semantic distance effects for pictures and weaker, less consistent effects for words. These results are inconsistent with strictly categorical views of semantic organization and are more consistent with representational frameworks in which semantic similarity is considered (see Frisby et al., 2023). While the hub-and-spoke account is not the only model compatible with semantic distance effects, it provides a comprehensive account that accommodates differences across picture and word formats, while simultaneously explaining a very wide range of other semantic results across multiple patient groups and healthy participants (Lambon Ralph et al., 2017). This study encourages further exploration of semantic architecture with the FPVS approach and bears promise for improving assessment of cognitive abilities in clinical populations such as patients with semantic dementia. Future studies should focus on assessing the effect of stimulation frequency on semantic distance effects depending on the stimulus input and on better understanding the interaction between picture and word stimuli in semantic discrimination processes, for example by presenting both stimulus types within the same visual sequence.

## Code and Data Availability Statement

Due to copyright reasons, pictures cannot be made freely available on a data repository but are available upon request to the corresponding author. All word stimuli are provided in Supplementary Material. Behavioural and EEG data are publicly available on an OSF repository: https://doi.org/10.17605/OSF.IO/BQC4X. No code was used as all data was analysed with the Letswave 6 toolbox in MATLAB.

## Acknowledgments

This research project was supported by an Early Career Researcher Grant (2022) from QUT Faculty of Health awarded to AV. MALR was supported by a Medical Research Council programme grant (MR/R023883/1) and intramural funding (MC_UU_00005/18).

## Open Access

For the purpose of open access, the UKRI-funded authors have applied a CC BY public copyright licence to any Author Accepted Manuscript version arising from this submission.

## Author contributions

Angelique Volfart: Conceptualization, Data curation, Formal analysis, Funding acquisition, Investigation, Methodology, Project administration, Resources, Visualization, Writing – original draft; Aliette Lochy: Writing – review & editing; Bruno Rossion: Conceptualization, Methodology, Supervision, Writing – review & editing; Matthew A. Lambon Ralph: Conceptualization, Methodology, Supervision, Writing – review & editing.

