## Supplementary Material for "Semantic distance differently modulates FPVS-EEG responses to words and pictures"

**List of all word stimuli included in each stimulus category**

| **Reference category** | **LD category** | **HD category** |
| --- | --- | --- |
| **Birds** | **Dogs** | **Furniture** |
| cuckoo  pelican  owl  swan  raven  eagle  goose  dove  duck  canary | labrador  corgi  bulldog  husky  poodle  spaniel  terrier  beagle  pug  alsatian | wardrobe  ottoman  mirror  bench  vase  cabinet  stool  lamp  rug  dresser |
|  | **Felines** | **Gardening tools** |
|  | ocelot  cheetah  lynx  leopard  cougar  jaguar  panther  tiger  lion  serval | axe  fork  hoe  mower  rake  scythe  shears  shovel  sickle  trowel |
|  | **Fish** | **Vehicles** |
|  | piranha  sturgeon  eel  tuna  sardine  mackerel  trout  carp  cod  catfish | airplane  tractor  canoe  sledge  ferry  wagon  barge  cart  tram  scooter |
|  | **Invertebrates** | **Kitchen utensils** |
|  | ant  mantis  slug  wasp  scorpion  snail  beetle  moth  bee  spider | spatula  strainer  sieve  scoop  funnel  scissors  masher  shaker  spoon  ladle |
|  | **Rodents** | **Musical instruments** |
|  | dormouse  gopher  raccoon  squirrel  shrew  rabbit  mouse  mole  rat  hamster | banjo  trombone  tuba  trumpet  cello  guitar  clarinet  harp  drum  flute |

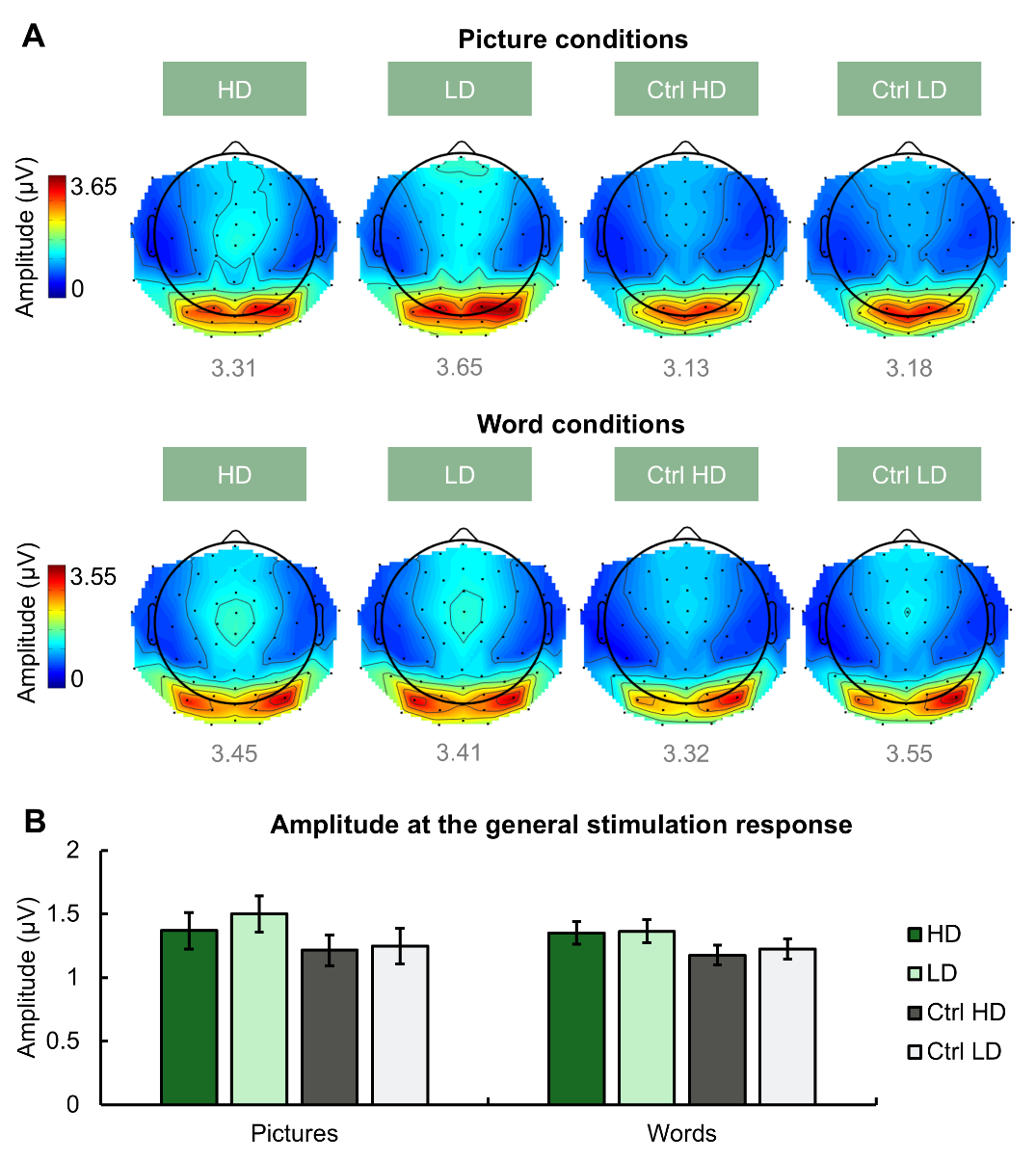

**Figure S1.** **A.** Topographical distribution of baseline-corrected amplitudes at the sum of 7 harmonics of the general stimulation frequency (grand-averaged data, n = 24 participants). The colour scale is adjusted to the maximal amplitude across picture and word conditions separately. The maximal amplitude in a given condition is shown in grey below each map. **B.** Baseline-corrected amplitudes at the sum of 7 harmonics of the general stimulation frequency across the whole scalp for picture and word conditions. Error bars represent ±1 standard error from the mean.
